# Methods to study xenografted human cancer in genetically diverse mice

**DOI:** 10.1101/2024.01.23.576906

**Authors:** Muneer G. Hasham, Jennifer K. Sargent, Mark A. Warner, Shawnna R. Farley, Brian R. Hoffmann, Timothy J. Stodola, Catherine J. Brunton, Steven C. Munger

## Abstract

Xenografting human cancer tissues into mice to test new cures against cancers is critical for understanding and treating the disease. However, only a few inbred strains of mice are used to study cancers, and derivatives of mainly one strain, mostly NOD/ShiLtJ, are used for therapy efficacy studies. As it has been demonstrated when human cancer cell lines or patient-derived tissues (PDX) are xenografted into mice, the neoplastic cells are human but the supporting cells that comprise the tumor (the stroma) are from the mouse. Therefore, results of studies of xenografted tissues are influenced by the host strain. We previously published that when the same neoplastic cells are xenografted into different mouse strains, the pattern of tumor growth, histology of the tumor, number of immune cells infiltrating the tumor, and types of circulating cytokines differ depending on the strain. Therefore, to better comprehend the behavior of cancer *in vivo*, one must xenograft multiple mouse strains. Here we describe and report a series of methods that we used to reveal the genes and proteins expressed when the same cancer cell line, MDA-MB-231, is xenografted in different hosts. First, using proteomic analysis, we show how to use the same cell line *in vivo* to reveal the protein changes in the neoplastic cell that help it adapt to its host. Then, we show how different hosts respond molecularly to the same cell line. We also find that using multiple strains can reveal a more suitable host than those traditionally used for a “difficult to xenograft” PDX. In addition, using complex trait genetics, we illustrate a feasible method for uncovering the alleles of the host that support tumor growth. Finally, we demonstrate that Diversity Outbred mice, the epitome of a model of mouse-strain genetic diversity, can be xenografted with human cell lines or PDX using 2-deoxy-D-glucose treatment.

## INTRODUCTION

Animal models have been used to dissect molecular, cellular, and physiological mechanisms of diseases (Abdolahi et al., 2022; Conn, 2017; Hidalgo et al., 2014; Robinson et al., 2019). After cardiovascular diseases, cancer is the leading cause of human fatality globally (de Magalhaes, 2013; Hamdi et al., 2021; Lin et al., 2021). To understand the basic progression of cancer *in vivo*, mouse models are used as a reliable and faithful platform for revealing underlying cellular and molecular mechanisms (Abdolahi et al., 2022; Liu et al., 2023; Zanella et al., 2022). Furthermore, mouse models are used to test the safety, efficacy, and bioavailability of any new therapy under consideration for human use (Cheon and Orsulic, 2011; Conn, 2017). There are two types of cancer mouse models for studies: a) spontaneous, or genetic, models, where a deleterious mutation is made in a mouse of a specific strain and these mice develop cancers to be studied (Chulpanova et al., 2020; Hill et al., 2021; Kersten et al., 2017); and b) models whereby human cancer tissues or cell lines are xenografted into immunodeficient mice, also known as patient-derived xenograft (PDX) or cell line-derived xenograft (CDX) models, respectively (Chen et al., 2022; Fujii et al., 2020; Hahn et al., 2021; Hidalgo et al., 2014; Jin et al., 2023; Klinghammer et al., 2017; Sargent et al., 2022; Shi et al., 2020; Uthamanthil et al., 2017; Yoshida, 2020; Zanella et al., 2022) These xenografts can either be subcutaneous or orthotopic depending on the context of the study.

Numerous cancer studies that use mouse models are performed in a single inbred strain. This is especially true in human xenograft mouse models, where one of the most common is the interleukin 2 receptor gamma-deficient NOD/ShiLtJ strain with either *Rag1* nullizygous or the severe combined immunodeficiency (*SCID*) mutation, abbreviated as NRG or NSG mice, respectively (Pearson et al., 2008; Shultz et al., 2005). The NOD/ShiLtJ strain is unique in that it expresses a protein, SIRP alpha, that allows the mice to tolerate the human immune system (Oronsky et al., 2020; Shultz et al., 2014).

Despite the utility of the above models, the lack of genetic diversity when using a single inbred strain limits the molecular, cellular, and physiological characteristics that can be revealed when studying a genetically complex disease such as cancer and narrows the scope of the etiological mechanisms that control the disease. For instance in GEMMs (Genetically Engineered Mouse Models), studies have shown differences among inbred Trp53-deficient strains in the length of time required for the development of malignancy: In the 129S1/SvImJ *Trp53-/-* and C57BL6/J *Trp53-/-* (p53 null) strains, mice develop neoplasias between 18-20 weeks (Donehower et al., 1992; Jacks et al., 1994; Olive et al., 2004). However, BALB/cJ *Trp53-/-* mice show neoplasias by 15 weeks, and in our studies of the tumor-prone SJL/J *Trp53-/-,* mice have neoplasias by 12 weeks (Branca et al., 2020; Kuperwasser et al., 2000). One would predict that a null mutation that completely removes the Trp53 protein (p53 in human) —a critical tumor suppressor and apoptosis regulator—would not be influenced by the strain background, since cell-cycle and apoptosis genes are conserved among mouse strains (Lozano and Levine, 2016). However, Trp53-deficient strains show not only a remarkable difference in cancer onset, but also in tumor-type spectrum; 129S/J *Trp53-/-* and C57BL6/J *Trp53-/-* mice develop primarily thymic lymphomas, whereas BALB/cJ *Trp53-/-* mice develop mammary tumors and SJL/J *Trp53- /-* mice develop testicular teratomas in addition to thymic lymphomas (Branca et al., 2020; Donehower et al., 1992; Jacks et al., 1994; Kuperwasser et al., 2000; Olive et al., 2004). Together, these results suggest that underlying genetic differences among the mouse strains influence the time and type of cancer development irrespective of the initiating mutation (Branca et al., 2020).

We recently published that the same homogenous human cancer cell line grows at different rates depending on the genetics of the lymphodeficient (*Rag1-/-*) host strain (Sargent et al., 2022). We also observed this phenomenon with the xenografted human breast cancer cell line MDA-MB-231, human chronic lymphocytic leukemia cell line MEC1, and mouse glioblastoma cell line GL261 (Sargent et al., 2022) For each of the three cell lines, tumors from that cell line exhibited differences in histological structure depending on the genetics of the host strain, even though the tumor-cell content itself was not impacted by host-strain genetics (Sargent et al., 2022) In addition, because myeloid cells are intact in the *Rag1-/-* host strains, our analysis showed strain-specific results for myeloid-population infiltration and composition in the tumors (Sargent et al., 2022) Finally, each *Rag1-/-* host strain exhibited a different chemokine and cytokine profile even though the number of cells xenografted was the same for each strain (Sargent et al., 2022). Together, these data suggest that different results may be obtained depending on which strain is used, whether studying a cellular or molecular mechanism, finding a biomarker, or targeting a pathway to test a new therapy (See editorial Dis Model Mech (2022) 15 (9)). Therefore, it is prudent to test more than one strain background when performing patient-derived xenografts in mouse model studies.

Here we describe methods for studying genetically diverse lymphodeficient and other genetically diverse mouse models “wildtype” (unmodified) mice to reveal genes and molecules that drive tumor growth. We show (i) the use of high-throughput tandem mass spectrometry of tumors to reveal protein-expression changes of the same cancer cell line and the host response in diverse strains; (ii) the use of xenograft models derived from F1 and F2 crosses to uncover quantitative trait loci (QTL) that determine tumor growth of a particular cell line; and (iii) a method to xenograft Diverse Outbred mice to study the growth of xenografted human cancer in a genetically diverse mouse population. Using these methods, one can begin to use genetically diverse mice to better approximate and define the cancer-host interactions that are crucial to cancer progression in the human population—results that can ultimately lead to new treatments and cures.

## MATERIALS AND METHODS

### Animal Use

All protocols and experiments that were performed were approved by The Jackson Laboratory (JAX) Institutional Animal Care and Use Committee (IACUC), and all regulations and accreditations are approved by the American Association for Accreditation of Laboratory Animal Care (AAALAC). Details on animal husbandry are described previously (Sargent et al., 2022).

### Cell Lines and Patient-Derived Xenografted Tissues

Cancer cell lines Hep G2 (HB-8065), HT29 (HTB-38), KG-1 (CCL-246), TOV-112D (HTB-161), and CCRF-SB were procured from ATCC; MDA-MB-231 was a kind gift from Dr. Karolina Palucka (JAX). All cell lines were grown in DMEM with glucose (Gibco), supplemented with 10% FBS (Gibco), sodium pyruvate (Gibco) and pen/strep/glutamine (Gibco) in 100mm tissue-culture dishes (ThermoFisher). PDX models TM00244, TM01245, and TM00089 were obtained from the JAX PDX Resource.

### Mice

All mice used in this study were obtained from The Jackson Laboratory. Inbred strains included 129S1/SvImJ (JR002448), 129S1/SvImJ *Rag1-/-* (129R, JR035322), A/J (JR000646), A/J *Rag1-/-* (A/JR, JR035069), BALB/cJ *Rag1-/-* (BAR, JR003145), C57BL/6J (JR000664), C57BL/6J *Rag1-/-* (B6R JR002216), CAST/EiJ (JR000928), NOD/ShiLtJ (JR001976), NOD/ShiLtJ *Rag1-/-* (NR, JR003729), NZO/HiltJ (JR002105), PWK/PhJ (JR003715), and WSB/EiJ (JR001145). For QTL mapping of tumor size in F2 intercross mice, C57BL/6J *Rag1-/-* (B6R) and NOD/ShiLtJ *Rag1-/-* (NR) parent strains were crossed, and the resulting F1 hybrid *Rag1-/-* progeny were then intercrossed to yield *Rag1-/-* F2 intercross progeny (n = 105 F2 mice) for QTL mapping. The F2 data is a combination of male B6R x female NR and female B6R x male NR.

For xenotransplantation in wildtype outbred mice treated with 2-Deoxy-D-glucose (2DG), 3-5-week old Diversity Outbred (DO) mice (J:DO, JR009376; n = 240 total DO mice) were obtained from The Jackson Laboratory. DO mice are produced by random matings in waves four times per year, with DO mice from generations 38-45 of outbreeding included in the current study.

### Xenografting Mice

Xenografting cells/tissues into mice was as described in Sargent et al. (Sargent et al., 2022). Inbred and DO mice were treated with 6g/L 2DG *ad libitum* in their drinking water one week before xenograft.

### Genotyping of F2 Intercross Mice

F2 intercross progeny are genetically unique and must be genotyped. F2 intercross mice (n = 105) were genotyped on the Mini Mouse Universal Genotyping Array (MiniMUGA, Transnetyx) at 2,340 SNP markers genome-wide that differ between the C57BL/6J and NOD/ShiLtJ strain backgrounds and passed a stringent quality filter (median gc score > 0.75). Pseudomarkers were imputed at 1-Mb intervals and inserted into the physical map using the *insert_pseudomarkers* function in the R package R/qtl2 (Broman et al., 2019).

### Quantitative Trait Locus (QTL) Mapping in F2 Intercross Mice

Following xenotransplantation of MDA-MB-231 cells into B6R x NR F2 intercross progeny, tumor size was measured with calipers at 2-, 3-, and 4-weeks post-transplantation. Tumor-size values were transformed to rank normal scores, and then genetic mapping was performed using a linear mixed model implemented by the *scan1* function in R/qtl2 package (Broman et al., 2019). Sex, cross direction, coat color, and batch were included as additive covariates, and the Leave One Chromosome Out (loco) option was used to correct for kinship. To estimate genome-wide significance of QTLs identified in the F2 analysis, genotypes were permuted 1,000 times while maintaining the sample-level relationship between the tumor-size phenotype and covariates; QTLs were deemed significant if they exceeded the LOD score corresponding to a permutation p-value < 0.05.

### Protein Extraction and Sample Preparation

Tumors in the study were snap frozen and stored at -80°C until samples were processed. To perform the protein extraction, a mixture of 50 mM HEPES, pH 8.2, containing 6M urea was added to each frozen sample at an equal weight-to-volume ratio, along with a pre-chilled 5-mm steel bead (QIAGEN). All samples were then placed in pre-chilled cassettes for the Tissue Lyser II and pulverized for at 30 1/s for three 1-minute oscillations. All pulverized samples were then waterbath-sonicated for 5 minutes (30 seconds on, 30 seconds off), followed by centrifugation at 21,000 x g at 4°C to remove heavy debris. Protein supernatants were then transferred to a new microfuge tube, diluted appropriately, and protein quantification was performed using the standard manufacturer microBCA protocol (Thermo, Cat.# 23235).

### Protein Digestion and Peptide Purification

After protein quantification, 100 µg of each sample were aliquoted and the urea was diluted 3- fold with 50 mM HEPES, pH 8.2 (brought <2M). Samples were reduced with 10 mM dithiothreitol at 42°C for 30 minutes on a ThermoMixer with agitation (500 rpm), alkylated with 15 mM iodoacetamide at room temperature for 20 minutes on a ThermoMixer with agitation (500 rpm), and trypsin-digested (Sequence Grade Modified; Promega) with a 1:100 trypsin:protein ratio at 37°C for 20 hours. Following the digest, all peptide samples were C18-purified using C18 MacroSpin Columns (Harvard Apparatus, Cat. #74-4101) according to the manufacturer’s protocol. Eluted purified peptides were dried with a vacuum centrifuge and stored at -20°C until TMT labeling was performed.

### Tandem Mass Tag (TMT) Pro Peptide Labeling

The dried peptide samples were reconstituted in 25 µL of 50 mM HEPES, pH 8.5) and quantified using the Quantitative Colorimetric Peptide Assay (Thermo, Cat. #23275) according to the manufacturer’s protocol. Samples were diluted to a concentration of 3.33 µg/µL in 50 mM HEPES, pH 8.5, and mixed on a ThermoMixer for 10 minutes at 25°C (500rpm). A pooled sample was made with equal amounts of all samples in the study for use as a carrier channel in each multiplex and for normalization purposes. While the peptide solutions mixed, 58.1 mM TMT stock solutions were created by adding 20 µL of dry acetonitrile for each 0.5 mg TMTpro reagent (Thermo, Cat. #A44520) and vortexing for 30 seconds. Immediately after reconstituting the TMT reagents, 4 µL of each TMT label (19.37 mM final concentration) was added to 12 µL (40 ug) of the appropriate peptide sample. All TMT tags were assigned to specific samples through a list randomization in Random.org. TMT peptide mixtures were then incubated for 1 hour at 25 °C with agitation on a ThermoMixer (500 rpm). Reactions were then quenched by adding hydroxylamine to a final concentration of 0.5% and incubating for 15 minutes at 25°C with agitation on a ThermoMixer (400 rpm). All samples were then combined in the respective TMT multiplex group and were acidified by adding 20 µL of 10% formic acid. Multiplexes were then snap-frozen and stored at -80°C overnight, followed by a drying step in a vacuum centrifuge the following day.

### Liquid Chromatography Tandem Mass Spectrometry Analysis (LC-MS/MS)

Dried multiplex samples were reconstituted in 50 µL of 0.1% TFA and zip-tipped using Millipore C18 zip-tips (Millipore, Cat. #ZTC18S096) according to the manufacturer protocol’s (Tadenev et al., 2023). Following zip-tip clean-up, samples were dried in a vacuum centrifuge and reconstituted in 20 µL of 98% H2O/2% ACN with 0.1% formic acid. Samples were then vortexed for 30 seconds, centrifuged for 30 seconds in a low-speed tabletop centrifuge, and transferred to a mass spec vial for the UltiMate 3000 autosampler. All LC-MS/MS was performed on a Thermo Eclipse Tribrid Orbitrap with a FAIMS coupled to an UltiMate 3000 nano-flow LC system in The Jackson Laboratory Mass Spectrometry and Protein Chemistry Service laboratory in Protein Sciences. The method duration was run at a flow rate of 300 nL/min with a 180-minute gradient of Buffer A (100% H2O with 0.1% formic acid) and Buffer B (100% acetonitrile with 0.1% formic acid) with a 15-minute equilibration using 98% A/2% B included. The exact gradient was 98% A/2% B from 0-2 minutes, 98% A/2% B at 2 minutes to 92.5% A/7.5% B at 15 minutes, 92.5% A/7.5% B at 15 minutes to 70% A/30% B at 145 minutes, 70% A/30% B at 145 minutes to 10% A/90% B at 155 minutes where it was held until 160 minutes, dropped to 98% A/2% B by 162 minutes, and was held there to equilibrate for 15 minutes. The TMT SP3 MS3 RTS method was used on the Eclipse Tribrid Orbitrap with global parameter settings including a default charge state of 2, expected peak width of 30 seconds, advanced peak determination, a static spray voltage of 2000V (Positive), FAIMS carrier gas at default settings, and an ion transfer tube temperature of 325°C. The instrument method consisted of three identical nodes with different FAIMS voltages of -40V, -55V, and -65V. Settings for precursor spectra detection (MS1) in each node included: cycle time = 1 second (each node), detector = Orbitrap, Orbitrap resolution = 120,000, scan range = 400-1600 m/z, RF lens % = 30, normalize AGC target (%) = 250, maximum inject time (ms) = auto, microscans = 1, data type = profile, polarity = positive, monoisotopic precursor selection = peptide, minimum intensity threshold = 5.0e3 (lower because of the FAIMS), charge states = 2-7, and a dynamic exclusion of a n = 1 for 60 seconds. Peptide fragment analysis (MS2) was performed in the ion trap and settings included: isolation window (m/z) = 0.7, collision energy (%) = 35 (fixed), activation type = CID, CID activation time = 10 ms, quadrupole isolation mode, ion trap scan rate = turbo, maximum inject time = 35 ms, and data type = centroid. Prior to data-dependent MS3 (ddMS3) analysis, real-time search was utilized with the canonical Uniprot Mus musculus (sp_tr_incl_isoforms) protein database with carbamidomethyl (+57.0215 Da on cysteine), TMTpro16plex (304.20171 on Kn), and oxidation of methionine (+15.9949). Additional parameters in the real-time search included maximum missed cleavages = 2, Xcorr threshold of 1, dCn threshold of 0.1, and a precursor ppm of 10. SP3 MS3 was performed in the Orbitrap and settings included SPS precursors = 20, isolation window = 0.7 m/z, activation type = HCD, HCD energy normalized at 45%, resolution = 60,000, scan range = 100-500 m/z, normalized AGC target = 500%, maximum injection time = 118 ms, and centroid data collection.

### LC-MS/MS Data Analysis

All of the Thermo Eclipse RAW mass spectrometry files were searched against the standard UniprotKB Mus musculus (sp_tr_incl_isoforms, TaxID=10090, v2021-02-04) and Homo Sapiens (SwissProt, TaxID=9606, v2020-11-09) databases using Sequest HT in Proteome Discoverer (version 2.5.0.400). Search parameters included trypsin digest, precursor mass tolerance of 20 ppm, fragment mass tolerance of 0.5 Da, statice carbamidomethyl on cysteines (+57.021 Da), static TMTpro modification on any N-terminus or lysine (+304.207 Da), and a variable oxidation of methionine (+15.995 Da). Other setting included a maximum number of missed cleavages = 2, minimum peptide length = 6, a maximum of 144 amino acids, a fragment mass tolerance = 0.6 Da and Percolator was used. In the Percolator node, the target/decoy selection = concatenated, with q-value validation, had a maximum delta Cn = 0.05, and a false discovery rate < 0.05 for all matches. The Minora node parameters were set to Thermo-recommended defaults, and data was filtered upon the target protein, and specific peptide maximum abundance/intensity values were extracted for each mouse model in the study using the Reporter Ions Quantifier Node for MS3 events.

### Other Software and Programs

Microsoft package for Mac (V.16) was used to generate the document, pie charts, tables, and figures. Prism for MacOS Version 9.5.0 (525) was used to generate graphs and statistical analysis. R statistical software (v 4.0.2) was used to generate QTL and allele effect plots. Pictures were obtained from JAX Creative.

## RESULTS

### The host strain determines protein expression of xenografted cancer cells

We previously showed, using multiple cancer-cell lines, that a given homogenous cancer-cell line xenografts at different rates depending on the genetic strain of the host mice, suggesting that the host provides critical support for the cancer cells to grow (Sargent et al., 2022). During our investigation of the cellular and molecular biology of the host, a question arose regarding whether the cancer cell changes its protein expression to suit the host. This is important to understand, especially if a drug target is chosen from tissue-culture results and the next step is to test the drug *in vivo* on cancer cells. Here we describe a method for selecting the best host to be xenografted with a cancer-cell line in order to target the protein of interest efficiently.

MDA-MB-231 cells, a triple negative breast cancer cell line, were xenografted subcutaneously in five strains as previously published Figure 1A, (Sargent et al., 2022). Collectively, these five strains provided the broadest spectrum with respect to *in vivo* tumor growth (Sargent et al., 2022). These strains are NOD/ShiLtJ *Rag1−/−* (NR), 129S1/SvImJ *Rag1-/-* (129R), BALB/cJ *Rag1−/−* (BAR), A/J *Rag1-/-* (A/JR), and C57BL/6J *Rag1−/−* (B6R). Female mice were chosen for this study so that the sex of the tumor matches the sex of the host. Other details about the host and xenograft procedure and data are published previously (Sargent et al., 2022).

**Figure 1:**
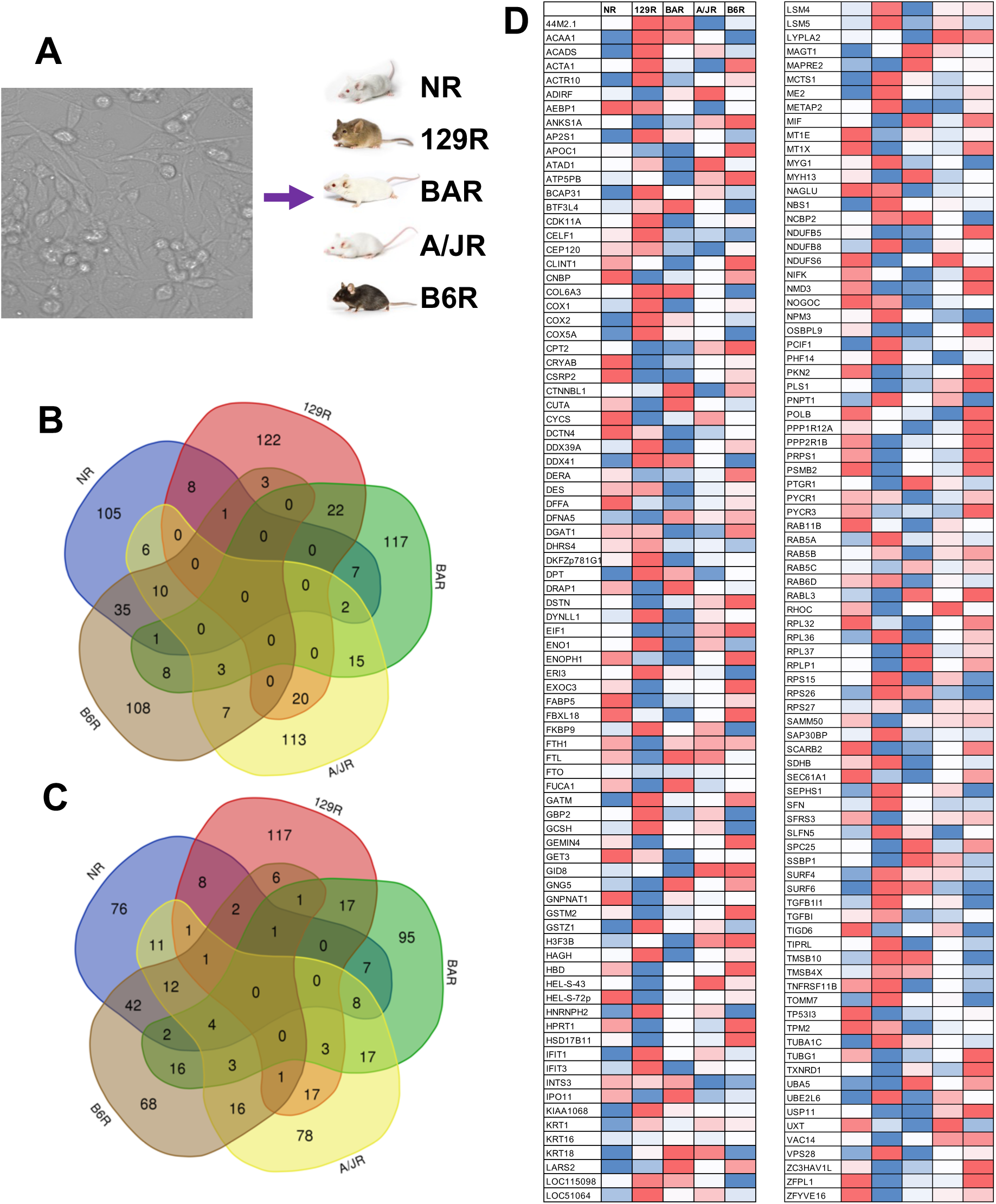
The host strain determines the protein expression of the xenografted cancer cells. (A) MDA-MB-231 cell line was xenografted in *Rag1-/-* 129S1/SvImJ (129R), A/J (A/JR), BALB/cJ (BAR), C57BL6/J (B6R), and NOD/ShiLtJ (NR) female mice (N=3 mice/strain). Mag = 40X. Tumors were harvested after 4 weeks, and (B) the 10% most abundant of all the detected human proteins (overexpressed), or (C) the 10% least abundant human proteins (repressed) were determined by mass spectrometry. Individual names of genes are listed in Supplementary Table 1. (D) List and expression of proteins produced by the human cancer cell in different mouse hosts as detected by mass spectrometry. 170 proteins differed by at least 2-fold when compared against at least one other mouse model host. Red to blue = low to high expression. Each biological sample was run through mass spectrometry three times.

As previously published, the tumors grew at different rates and sizes (Sargent et al., 2022). We chose the largest, the median, and the smallest tumor size within each strain to perform tandem mass spectrometry-based proteomics analysis to determine the changes in protein expression in both the xenografted cancer-cell line and in the host cells that compose the micro-environment. Proteins were extracted from the tissues, processed, TMT-labeled, multiplexed, and run via LC-MS/MS as described in Materials and Methods. The protein-expression data had shown there were 1,759 detected human proteins identified, relative to a total of around 4 thousand proteins contributed by the host mouse models (Supplementary Table I). Scaled protein abundances were generated in Proteome Discoverer based on the normalized intensity values across all samples for comparison. This abundance was the median of the abundances in the afore-described three sample types (largest, median, smallest tumor size) with percent confidence values (%CV). For each strain, the most to the least abundant human proteins detected were sorted (Supplementary Table II), and the 10% most abundant and 10% least abundant human proteins in the dataset were determined. These 175 human proteins from each strain were then compared for similarity using an online Venn diagram software tool (http://bioinformatics.psb.ugent.be/webtools/Venn/). The results show that for either the least abundant (Figure 1B) or most abundant (Figure 1C) human proteins in each strain, no proteins overlapped between all strains. However, the results also show that there are proteins shared between various host strains. This suggests that the human cancer cell itself changes its protein expression depending on the host or the host compensates differently. The full list and functions of the proteins shared/not shared between host strains can be seen in Supplementary Tables III and IV).

Our results above raise an issue about choosing a mouse strain in which to xenograft a cell line to develop the xenografted mouse as a model for testing a compound or even for accurately studying the biology of a cancer. To illustrate this issue, we filtered the list of 1,759 detected human proteins to the 171 proteins that showed at least a two-fold difference in expression in one strain compared to the other four strains (Figure 1D, Supplemental Table V). The data definitively shows the importance of carefully evaluating a strain to be used for xenograft before testing a compound *in vivo* or studying the biology of a cancer. For instance, the protein CDK11A is expressed most abundantly by MDA-MB-231 cells when xenografted in BAR mice compared to the other four strains. Therefore, when testing CDK11A inhibitors such as the FDA- and EMA-approved drug Crizotinib, which is used for breast cancer, BAR would be a better model than others, such as the two NOD-derived strains that are commonly used as the standard for drug efficacy studies—NSG and NRG mice (Ayoub et al., 2017; Whittaker et al., 2017). Altogether, these results suggest that the genetics of the host must be considered before using a mouse model to test a compound/therapy or study the biology of a cancer, because not all proteins of the xenografted cells are expressed in the same quantities in all hosts.

### Genetically diverse *Rag1-/-* hosts reveal proteins that determine a cancer phenotype

Previously, we showed that engraftment of MDA-MB-231 cells into B6R host mice resulted in collagen accumulation in the tumors, while engraftment of these cells into NR, 129R, or BAR mice did not result in that phenotype (Sargent et al., 2022). Here we illustrate a method using tandem mass spectrometry analysis to reveal proteins that determine a phenotype. Identification of such proteins can enable either targeting the protein(s) to eliminate the phenotype in the host strain exhibiting the phenotype or increasing the abundance of the protein(s) to enhance the phenotype. In our previously published result referred to above, the mass spectrometry data shows not only the human proteins but also the mouse proteins, since a tumor is composed of both human neoplastic cells and mouse host cells. Therefore, using the same mass spectrometry data generated in our previous study above, we could computationally separate the human proteins, which were used for the analysis above, and the mouse proteins, which we used for this analysis. This separation was achievable because protein annotation differs between human and mouse.

The host protein abundance of the NR, 129R, and BAR strains was compared, separately, to the protein abundance of B6R, the strain exhibiting collagen deposits (Supplemental Table VI). Then, the following protein abundance ratios were obtained: NR:B6R, 129R:B6R, and BAR:B6R. The ratios were then sorted, and all the proteins that were expressed 2-fold or lower in NR, 129R, or BAR compared to B6R (ratio of <0.51, Supplemental Table VI in bold) were compared to each other as a Venn diagram (Figure 2A). Four proteins—CES1d, RB3, SERPIN1d, and SERPINa3k—were found in lower abundance in all three strains that lacked collagen deposits, and these proteins have been shown to be essential in collagen deposition and related biology (Figure 2A-B) (Cox, 2021; Law et al., 2006; Tsukui et al., 2020; Wang et al., 2018).

**Figure 2:**
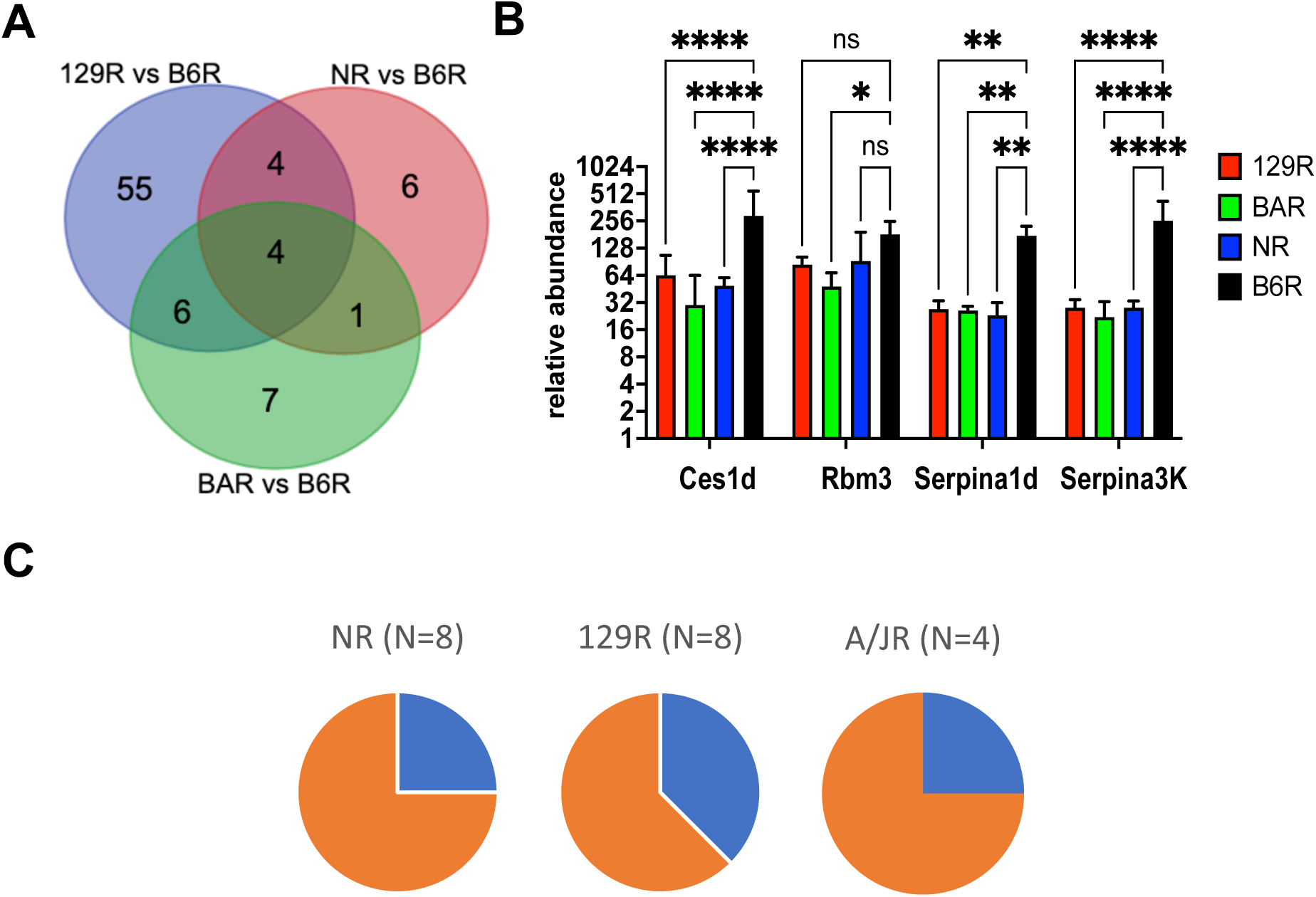
Analyzing xenografted host protein expression from multiple *Rag1-/-* strains reveals proteins that determine a phenotype. (A) MDA-MB-231 cell line engrafted into B6R and A/JR mice previously showed high levels of collagen deposition compared to the same cell line engrafted in 129R, BAR, or NR mice. (B) When relative protein expression was compared between B6R (high collagen) to 129, BAR, or NR (low collagen), four essential collagen- associated host genes were upregulated in B6R mice: Serpina1, Serpina3k, Rbm3, and Ces1d. Graphs showing relative abundance of the four aforementioned genes. Six biological replicates per strain. Error bars are 95% CV of each sample analyzed at least 3 times. (C) Xenografting with PDX has a higher probability of success in genetically diverse *Rag1-/-* hosts than in NSG mice. For example, PDX J000101173 (invasive ductal carcinoma Stage III) xenografted better in three out of four *Rag1-/-* mice than in NSG mice.

As illustrated above, xenografting different strains and then using the comparative protein data can reveal key proteins that control the disease phenotype. In fact, these *Rag1-/-* mice can potentially be more receptive to xenografts compared to the well-known NSG mice. For instance, an invasive ductal carcinoma stage III patient-derived xenograft (PDX) (J000101173) is recalcitrant in NSG mice, giving us one positive xenograft in 11 attempts, whereas the same PDX gave us two out of eight in NR mice, three out of eight in 129R mice, and one out of four in A/JR mice (Figure 2C). The B6R strain, which has shown poor growth for multiple types of cancers, also shows no xenograft capability with this PDX (Figure 2C). Together, these results suggest that expanding the choice of host strain to multiple genetically diverse hosts could not only provide a tool for revealing the proteins that control the phenotype, but also potentially improve the outcome of xenografts for recalcitrant human tissue. It has been shown that several cancers have less than 50% chance of xenografting (Goto, 2020). Perhaps xenografting different hosts may be the key to success to rescuing recalcitrant human tissues.

### QTL mapping identifies genetic modifiers of tumor growth in the NR and B6R strains

Based on the observed variability in tumor growth and protein abundances among the *Rag1* knockout strain backgrounds, we reasoned that genetic variation may underlie these differences. To maximize the likelihood of identifying genetic modifiers of tumor growth, we focused our genetic analysis on the B6R and NR strains, as these strains were previously shown to differ significantly in tumor growth rates following xenotransplantation with MDA-MB-231 breast-cancer cells (Sargent et al., 2022). We see similar strain differences in this study; four weeks post-xenograft with the same starting amount of tissue (20 mm^3^), tumors in NR mice were significantly larger than those in B6R mice (Figure 3A). Tumor growth in F1 hybrid progeny from the cross of NR to B6R mice is more variable than that in either parent strain, suggesting that the observed parent-strain differences are not due to a single dominant modifier allele. Next, we intercrossed the F1 hybrid mice to create a population of F2 mice (n = 105). Each F2 mouse is a genetically unique combination of the two parent strain genomes; as expected, we measured highly variable tumor growth in these diverse mice (Figure 3A). Some growth differences in the F2 mice appeared to be linked to sex (Supplementary Figure 1A) and coat color (Supplementary Figure 1B), suggesting that genetic variants influencing tumor growth may be found on the sex chromosomes or linked to autosomal coat-color alleles (i.e., Tyrosinase *Tyr*, Nonagouti *a*).

**Figure 3:**
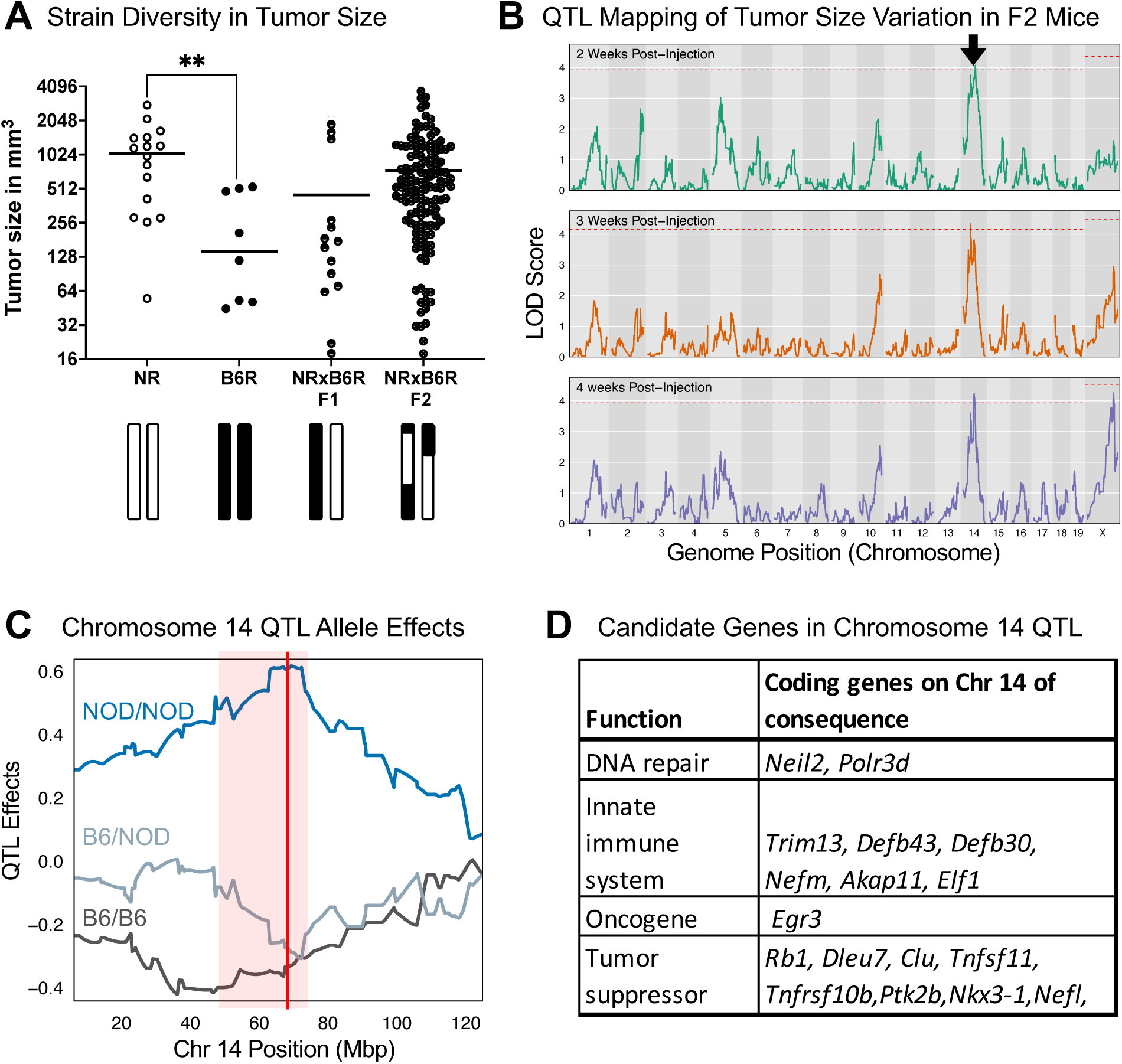
Quantitative trait locus (QTL) mapping reveals a genetic modifier of tumor growth on Chromosome 14. (A) Four-week tumor growth of MDA-MB-231 cells in the best host (NR), the worst host (B6R), and the first (F1) and second (F2) filial generation hosts. (B) QTL plots showing the Logarithm of Odds (LOD) significance score for tumor growth at 2, 3, and 4 weeks post-injection. One region on Chromosome (Chr) 14 denoted by the black arrow surpasses the genome-wide LOD significance threshold (LOD > 4.06). (C) Strain allele effects at the Chr 14 QTL. Tumors in F2 mice homozygous for the NOD allele (blue line) in this region are larger than those in individuals that are homozygous B6 or heterozygous NOD/B6. QTL peak position and 90% Bayesian confidence interval are denoted by a red line and pink box, respectively. (D) Genes in the Chr 14 QTL that are strong candidates for being the causal gene(s) based on their established roles in cancer.

To identify genetic factors responsible for these strain differences in tumor growth, we performed quantitative trait locus (QTL) mapping on the F2 mice, since the F1 mice are all heterozygous on all loci. First, we genotyped each F2 mouse as well as a subset of B6R, NR, F1 control samples at 2,340 SNP markers genome-wide that differ between these two strains (MiniMUGA genotyping array, Transnetyx; see Materials and Methods). Next, we applied a linear mixed model using the R package R/qtl2 to identify regions of the genome (QTLs) where genotype differences among the F2 mice were linked to quantitative tumor growth differences at 2, 3, and/or 4 weeks after xenotransplantation (Figure 3B) (Broman et al., 2019). We identified a significant QTL on Chr 14 centered at 67.0846 Mb (90% Bayesian confidence interval 47.2 – 73.5 Mb; permutation p-value = 0.028) that was associated with tumor growth as early as 2 weeks post-transplantation (denoted by arrow in Figure 3B). F2 mice that are homozygous for the NOD allele at this locus exhibit larger tumor size on average than those that are heterozygous or homozygous B6 in the same region, suggesting that a single B6 allele in this region is sufficient to protect against tumor growth (Figure 3C). We found several plausible candidate genes among the 361 protein-coding genes in this critical region that are known to be associated with cancer susceptibility and/or tumor growth and may be causal for the observed strain differences in this analysis (Figure 3D). Of note are *Egr3* and *Rb1* tumor suppressor genes, which are found close to the QTL peak and together have >500 citations associating them with cancer growth (for example: (Chinnam and Goodrich, 2011; Dyson, 2016; Shin et al., 2020; Yao et al., 2022). Finally, we identified a suggestive tumor-size QTL on the X Chromosome that is evident at 4 weeks post-transplantation but not at earlier timepoints (peak position 145.3162 Mb, 90% Bayesian confidence interval 120.4 – 152.4 Mb; permutation p-value = 0.1; Figure 3B). In contrast to strain effects observed for the Chr 14 QTL, F2 mice that are homozygous B6 in this region of Chr X have, on average, larger tumors than mice homozygous for the NOD allele (Supplementary Figure 2). Interestingly, tumors in heterozygous mice at this locus are smaller than those in either of the two homozygous parental strains, suggesting that heterozygosity in this region may confer resistance to tumor growth. Overall, results of this analysis reveal loci associated with tumor-growth variation; highlight that each inbred parental-strain (B6R, NR) genome harbors multiple alleles that are either beneficial or detrimental to tumor growth; and suggest that specific combinations of inbred-strain genomes that, together, align pro-growth alleles may yield a mouse stroma with a tumor microenvironment that is especially receptive to transplanted human tumors.

### Immunosuppressant 2-deoxy-D-glucose aids xenografts

The absence/disruption of the *Rag1* gene eliminates B- and T- lymphocytes in the mouse, resulting in lymphodeficient mice and consequently allowing acceptance of human xenografts (Oettinger, 1992; Shultz et al., 2014; Sobacchi et al., 2006; Zhu and Roth, 1996). While the generation of several *Rag1-/-* strains is feasible, it is costly and time-consuming to generate enough F2 mice to map QTLs. Therefore, new methods are necessary to allow xenografts to be tolerated in “wildtype” (genetically unmodified) mouse models. While working with glycolysis-resistant tissues, we discovered that treatment with the inhibitor 2-deoxy-D-glucose (2DG) improved xenografts in immunodeficient NSG mice (Wilson et al., 2019). The immunosuppressant 2DG actively competes with glucose for hexokinases (Pajak et al., 2019; Warburg, 1956). Glucose is required by the immune system to respond to antigens expediently, as most immune cells do not carry a fat reserve (Ganeshan and Chawla, 2014). It turns out that while numerous cancers are sensitive to 2DG, some cancers are glycolysis-resistant and are unaffected by 2DG (Wilson et al., 2019). When we xenografted NSG mice after they were pre-treated with 6g/L 2DG *ad libitum* in their drinking water for one week, a glycolysis-resistant cancer, i.e., the lung squamous cell carcinoma PDX TM00244, grew significantly better in these 2DG-treated treated mice than in mice pre-treated with vehicle (water) (Figure 4A). Use of 2DG has multiple advantages: It is a relatively inexpensive immunosuppressant; it is administered *ad libitum*; and the mice do not show adverse effects such as weight loss (Figure 4B).

**Figure 4:**
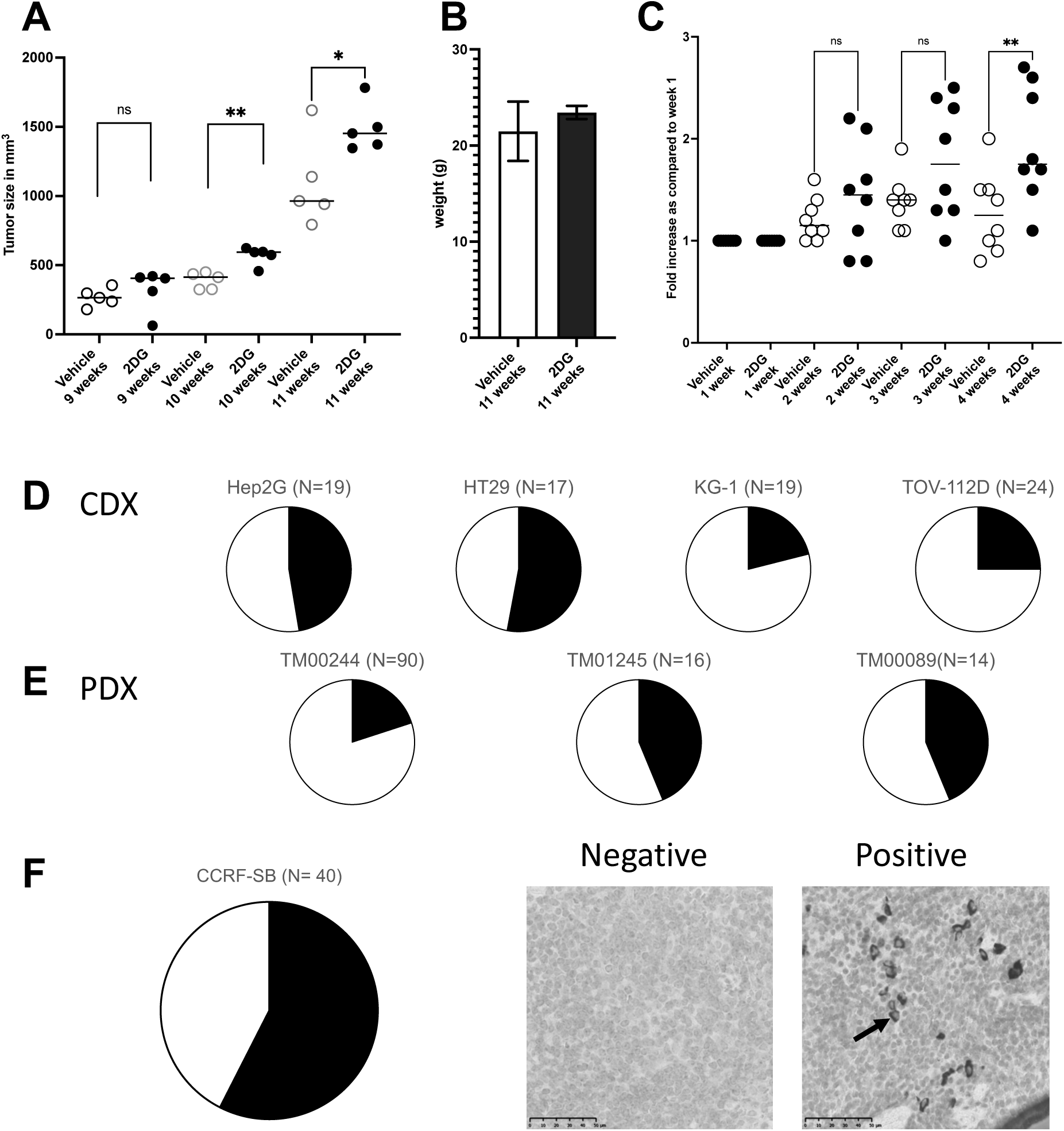
2-deoxy-D-glucose (2DG) is a unique inhibitor that improves the growth of xenografted tissues or cells in a diverse wildtype mouse population. (A) Graph showing the improvement in growth of a glycolysis-resistant tumor, TM00244, in NSG mice treated with 2DG (black circles) verses vehicle (white circles). (B) Graph showing no differences in weight between 2DG-treated mice and vehicle-treated mice. (C) Individual inbred mice (eight strains), viewed as a population, show improved xenograft growth of the MDA-MB-231 cell line when treated with 2DG. Black circles, 2DG-treated; white circles, vehicle-treated. (D, E) Pie charts showing the proportion of 2DG-treated DO mice that can accept CDX or PDX. The number of DO mice used for each cancer type is shown in the parentheses. Any tumor >100mm^3^ as measured by calipers was deemed positive (black shaded part) (F) As a leukemia, CCRF-SB, was measured by the presence (positive) or absence (negative) of infiltrating human CD19+ cells in the spleen. Pie charts show the fraction of positive (black) versus negative (white) spleens from 2DG treated DO mice.

To test the extent to which 2DG could be used as an immunosuppressant and allow xenografts, we obtained one male and one female mouse from each of the eight parental founders inbred strains of the Collaborative Cross (CC) and Diversity Outbred (DO) mouse populations. Together, these eight founder strains segregate nearly 50 million single nucleotide variants (SNVs) and millions of short indels and larger structural variants (Chesler et al., 2016; Churchill et al., 2012; Lilue et al., 2018). We pretreated the sixteen mice *ad libitum* with 6g/L 2DG in their drinking water. A similar cohort was obtained and were untreated (vehicle). After one week, the MDA-MB-231 breast-cancer cell line, which is glycosensitive, was xenografted and tumor size was monitored. As populations, the diverse inbred strains treated with 2DG showed significantly better tumor growth compared to the untreated mice (Supplementary Figure 3). The differences in tumor growth observed at 4 weeks suggest that 2DG could be a viable method of immunosuppressing wild-type inbred mice to accept xenografts. Interestingly, some strains showed better acceptance than others, which appears to be due to genetic background differences as all other factors are controlled for. (Supplementary Figure 3).

### Xenografting immunocompetent Diversity Outbred (DO) mice could improve novel allele identification

The *Rag1-/-* inbred strains highlight how genetic diversity in the host stroma can impact neoplastic growth, and our F2 mapping study points to two genomic loci—one on Chr 14 and one on Chr X—that may underlie the observed variability in tumor growth between the NR and B6R strains. However, the need to include the homozygous *Rag1* knockout allele limits the number of strains, due to costs, that we can study, thus hindering our ability to identify novel alleles and their genetic interactions that influence tumor growth. We reasoned that 2DG treatment may enable successful xenotransplantation of difficult tumors and/or tumor cell lines in a subset of wildtype DO mice. DO mice are derived from the eight CC/DO founder strains and maintained by random outcrossing, resulting in a population of genetically unique individuals that segregate high genetic variation and are optimized for genetic-mapping studies that approximate genetic diversity in the human population.

To illustrate the capability of xenografting DO mice and thus establish their potential as a resource for future genetic-mapping studies, we chose five human cancer-cell lines (CDX) and three human patient-derived tissues (PDX). The criterion for selecting these cell lines is that the cells/tissues had to be either 2DG-tolerant or -resistant. The basis for this criterion is that DO mice are immunocompetent, and thus xenografting human tissues/cells will lead to rejection unless the immune system is suppressed. As illustrated above, 2DG immunosuppresses different inbred strains of mice such that they accept xenografts; therefore, we chose to use 2DG to suppress the immune system of the DO mice. However, we and others have shown that 2DG is a chemotherapy for cancers, and therefore the CDX and PDX chosen must be 2DG-resistant or at least tolerant. Resistance or tolerance was tested by xenografting the CDX or PDX in NSG mice treated with 2DG and measuring tumor growth over time (data not shown).

Cell lines chosen included Hep G2 (hepatoma), HT29 (colorectal adenocarcinoma), KG-1 (acute myelogenous leukemia), TOV-112D (ovarian endometrioid adenocarcinoma), TM00244 (lung squamous carcinoma), TM01245 (lung adenocarcinoma), and TM00089 (invasive breast carcinoma), which were xenografted subcutaneously in DO mice; and CCRF-SB (B-lymphocyte acute lymphoblastic leukemia), which was xenografted in DO mice via the tail vein. For all CDX and PDX tested, a subset of 2DG-treated DO animals were able to grow tumors (Figure 4D-E), with some cell models showing higher (HT29) or lower (KG-1) success rates among DO mice. Together, these results are consistent with our expectation that some combinations of alleles in individual DO mice would allow tumors to grow while other combinations would inhibit growth. For CCRF-SB cells, since they had to be grown orthotopically, the spleen was used as an indicator of cancer cell growth. Spleens were fixed, sectioned and stained for human CD45+ cells (white blood-cell marker), and scored for the intensity of infiltration. We scored the spleen sections as either negative or positive for CD45+ cells (Figure 4F). We observed human CD45+ cells indicating tumor growth in over half of 2DG-treated DO individuals (Figure 4F). Given the observed data showing growth of all PDX and CDX models in at least a subset of DO animals, we expect that future QTL mapping studies in DO mice will be possible and likely to succeed in revealing additional tumor growth-associated genes and genetic variants.

## DISCUSSION

We recently showed that xenografting the same human cancer cells in genetically diverse mouse strains yielded different tumor-growth kinetics across these strains. (Sargent et al., 2022). Furthermore, in this study we show that, despite starting with the same human cancer cells, the cellular and molecular signatures of the resulting tumors are diverse and strain-dependent. This suggests that the host stroma that supports the cancer cells plays a major role in the development and molecular signature of cancer. It has been suggested that one could view the neoplastic cell as a ‘seed’ and the micro-environment as the ‘soil’, whereby a good seed is unable to develop and grow in bad soil (in bad soil, and bad soil is unable support growth of a good seed. This idea, originally coined in 1889 by Paget on the nature of cancer metastasis, can now be adapted to local tumor growth as shown in our previous and this study; the microenvironment is a primary contributor of neoplastic growth (Paget, 1989; Sargent et al., 2022). Understanding the biological interaction of the host micro-environment and the neoplastic cell is essential in uncovering novel molecular pathways to target, providing improved treatments and cures for humans.

Studying the tumor as a whole from patients *ex vivo* is limited. Each human being is exposed to countless variables that could influence the tumor microenvironment, and it would be difficult to draw biological conclusions unless many patients are surveyed. In addition, since the cancerous cells and the cells that compose the micro-environment come from the same individual, it is difficult to molecularly separate the genetics and subsequently expression, whether mRNA or protein, of the cancer cell from the cells of the microenvironment. This makes mapping host genetic factors in human populations exceedingly difficult. The mouse model allows us to interrogate growth of a single tumor genome in the context of multiple genetically diverse stroma, and, furthermore, enables us to identify the specific genes and genetic variants driving this stromal variation.

Previously we published a resource article generating a series of genetically diverse lymphodeficient mice that can accept human cancer cells (ref). Here we illustrate the use of the lymphodeficient (*Rag1*-/-) and other more complex genetically diverse (DO) mice to uncover genetic and molecular mechanisms that allow the tumor microenvironment to support the neoplastic cells and allow the molecular response of the neoplastic cells to adapt in genetically diverse microenvironments.

Our first method of protein profiling shows that one can observe the molecular changes of a cancer cell of interest to determine the type of host *in vivo* molecular response that is initiated. This information is essential to know when choosing a suitable host in which to grow tumors to test a drug of interest. We observe that protein expression of a given cancer cell differs in different hosts; thus, using the wrong host may yield inaccurate efficacy data. In addition to the aforementioned example, the breast-cancer cell tested, MDA-MB-231, expresses TP53I3 protein more highly in the 129S1/SvImJ background (129R) than in the NOD/ShiLtJ background (Figure 1D; the latter strain is NR mice, which have the same background as that of NSG or NRG standard mice). Therefore, testing a drug that targets TP53I3 in NSG-background mice may not yield discernable results, as expression in this genetic background is low; the appropriate strain to use would be 129R, in this case. In Figure 1D, we show numerous examples of proteins that exhibit vastly different expression in different strains. We chose TP53I3 as an example because it is regulated by the p53 protein, which is known to be a major tumor suppressor. An argument can be made that inter-strain expression differences greater than two-fold are seen in only 10% of the total detected proteins of the cancer cell in our study. However, one cannot predict the degree to which expression of a given tumor or microenvironment gene or protein will differ among different strains. Therefore, it is cost-effective to test one’s cancer cell line or PDX of interest in a range of genetically diverse lymphodeficient strains before launching efficacy studies.

To perform the aforementioned proteomic experiment/analysis, one must use mouse models; performing such experiment/analysis directly in human tumors will not allow separation of the proteins that arise from the stroma from those that arise from the neoplastic cells, since proteins from both sources will be annotated in the same way. Since many human proteins differ in sequence from their mouse orthologs, one can distinguish between the stroma proteins, which come from the mouse host, from the human neoplastic cells in proteomics datasets. Therefore, our method can clearly differentiate between proteins from the stroma and proteins from neoplastic cells. This ability will in turn enable investigators to investigate proteins made by the host without contamination by the neoplastic cells. One feature we previously showed is that tumors from B6R mice contained large deposits of collagen and were smaller than the tumors found in NR, 129R, or BAR mice (Sargent et al., 2022). When we compared the most abundant host proteins expressed by the largest tumors from the hosts NR, 129R, and BAR versus B6R, four proteins were shared among all the hosts, and each protein regulates collagen deposition, as aforemetioned. Thus, using these diverse genetic models, one can reveal proteins that are shared by different strains and that lead to a particular phenotype of interest. In fact, although NSG mice are a great model, our genetically diverse mice may be capable of functioning as better hosts for stubborn cell lines and PDXs.

Importantly, in this study we show that identification of host genes with a major impact on tumor growth can be accomplished not only via molecular analysis of the human tumor and mouse stroma but also via genetic mapping. First, we observed that growth of MDA-MB-231 cells was most prolific in NR mice and least prolific in B6R. We then mated the two strains and subsequently intercrossed the F1 progeny to generate a mapping population of genetically unique F2 mice. For genetic mapping, we recommend a minimum of 100 F2 mice; in the current study, 105 F2 mice were sufficient to map a significant tumor-growth QTL on Chr 14 and a suggestive QTL on the X Chromosome. While the purpose of this study was to demonstrate the power of this mapping platform, it will be interesting to determine if mapping studies of different cancer cell lines in this or similar F2 populations will identify the same genomic loci or novel QTLs associated specifically with the particular tumor under investigation. Finally, it should be noted that testing as few as 20 F2 mice is sufficient to estimate the range of tumor growth in the population, and that this knowledge can be used in power calculations to estimate the sample size needed to map significant QTLs. Given the costs associated with genotyping intercross mice, we recommend that investigators engrafting a new cancer model should perform such a pilot study in a subset of F1 hybrid and F2 intercross progeny prior to expanding to a much larger population for genetic mapping.

The immune system is a critical element in fighting cancer (Weinberg, 2014). Indeed, this is why NSG and NRG mice are ideal hosts for studying immunotherapies; not only do they lack an adaptive immune system, but they also have a severely compromised innate immune system, and their mutual NOD background inbred strain carries the *Sirp* allele that tolerates engraftment of the human immune system (Oronsky et al., 2020; Shultz et al., 2014). The myeloid components of these *Rag1-/-* strains of mice are intact, and therefore while these mice may or may not be capable of hosting the human immune system, they could be used to study the ‘native’ myeloid immune response to tumors in the absence of the adaptive immune system. We showed that, in the *Rag1-/-* strains examined in the present study, with the exception of NOD/ShiLtJ, tumor size is positively correlated with the number of myeloid cells in the tumor (Sargent et al., 2022). The mechanism underlying this phenomenon still needs to be studied.

An alternative strategy to using genetically engineered strains like *Rag1-/-* for tumor engraftment is to treat wildtype animals with the immunosuppressant 2DG. This method enables one to expand their analysis of host factors to highly genetically diverse populations like the Diversity Outbred and Collaborative Cross strains that were specifically designed to provide optimal power for genetic mapping studies. In this study, we showed that xenotransplantation of several PDX models and CDX cell lines was successful in at least a subset of 2DG-treated DO mice, indicating that genetic mapping is feasible and will likely be successful in larger populations of DO mice.

In summary, here we explore various genetic and proteomic methods to study xenografted cancer in genetically diverse mice. As we have shown above, these methods reveal the alleles and proteins that change the phenotype of a given cancer cell depending on the genetic background of the host. We speculate that these powerful techniques will be utilized to further understand the biology of, and find new therapies for, cancer.

## Author contributions

Conceptualization: M.G.H.; Methodology: M.G.H., J.K.S., M.A.W., S.R.F., C.B., T.S., S.C.M, B.R.H.; Formal analysis: M.G.H., B.R.H., S.C.M; Writing - original draft: M.G.H., B.R.H., S.C.M; Reviewing and editing: M.G.H., J.K.S., M.A.W., S.R.F., C.B., B.R.H., S.C.M.; Supervision: M.G.H., S.C.M, B.R.H.

## Acknowledgements

We want to acknowledge the following persons and Scientific Services at The Jackson Laboratory for their contribution to this body of work: Elaine Bechtel, Nick Gott and Histology Department; Dorothy Ahlf-Wheatcraft and Mass Spectrometry and Protein Chemistry Service within Protein Sciences; and Ali Jones and the Xenografting and Live Imaging Service (formerly the PDX R&D Core). We thank JAX Creative for the mouse images in Figure 1. We also thank Stephen Sampson for manuscript editing. MDA-MB-231 cells were a kind gift from Dr. Karolina Palucka (The Jackson Laboratory). Open Access funding was provided by The Jackson Laboratory.

