## Supplemental Figures for "Methods to study xenografted human cancer in genetically diverse mice"

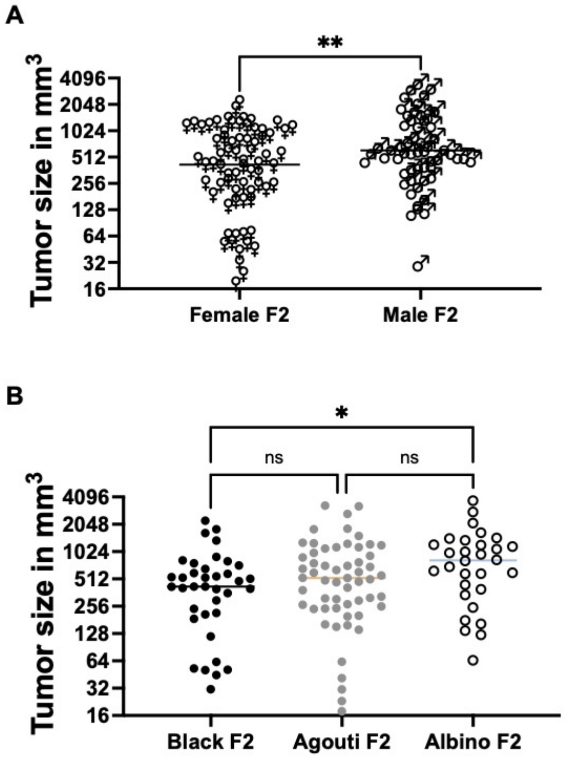


**Supplemental Figure 1. Tumor growth in F2 mice is correlated with sex and coat color.** (A) Tumor size is significantly greater in F2 males as a group compared to females ($p < 0.01$). (B) Tumor size is significantly greater in albino F2 mice as a group compared to black F2 mice ($p < 0.05$).

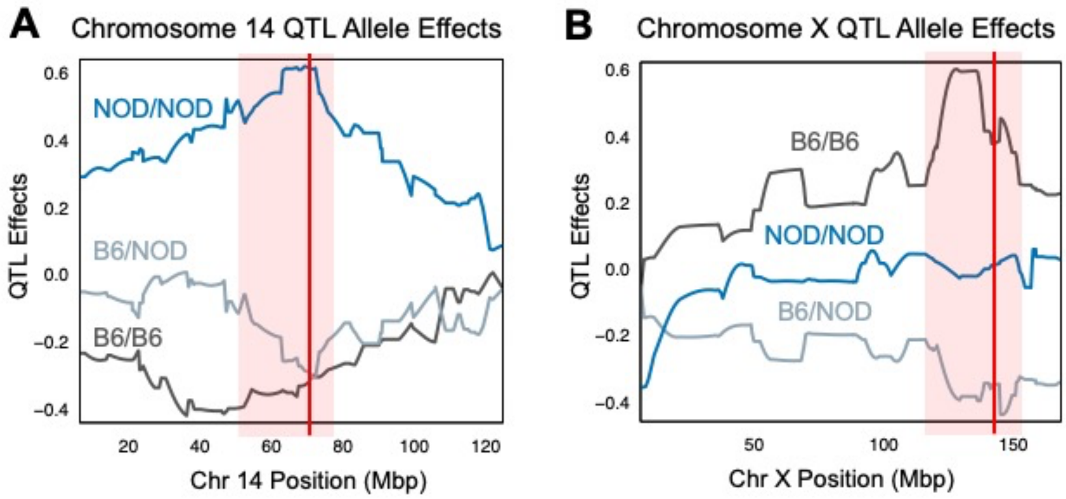


**Supplemental Figure 2. Strain allele effects at tumor size QTLs.** (A) Strain allele effects at significant Chromosome 14 tumor growth QTL shows that F2 mice homozygous for the NOD allele at this locus exhibit larger tumors than those F2 mice that are homozygous B6/B6 or heterozygous B6/NOD. (B) Strain allele effects at the suggestive Chromosome X tumor growth QTL shows that F2 mice homozygous for the B6 allele at this locus exhibit larger tumors compared to F2 mice homozygous NOD/NOD or heterozygous B6/NOD.

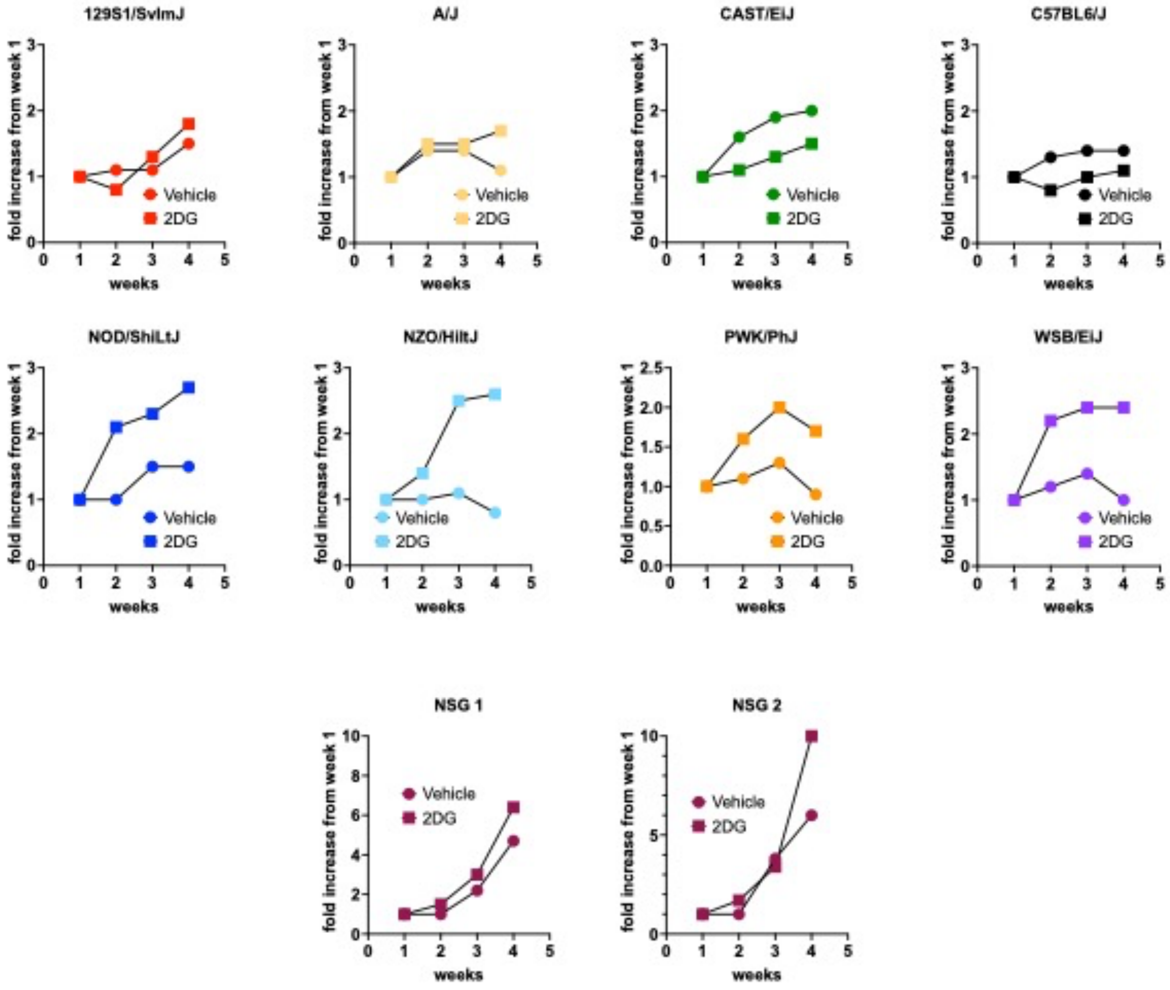


**Supplemental Figure 3. Effect of 2DG treatment on growth in individual strains with MDA-MB-231.** Inbred strains differ in their response to 2DG treatment on tumor growth, with some strains including 129S1/SvImJ showing little effect from 2DG, while others including NZO/HiltJ exhibit increased tumor growth in response to 2DG.
